# BUsmear: A Low-Cost 3D-Printed Automated Device for Blood Smear Preparation

**DOI:** 10.64898/2026.08.03.742560

**Authors:** Tuna Pesen, M. Talha Karasoy, Beyza Ceren Eren, Bora Akgun

## Abstract

Uniform, reproducible blood smears are critical for reliable hematological evaluation. Manual smear preparation, however, is user-dependent and introduces variability that limits quantitative microscopy. Here we developed BUsmear, a low-cost, 3D-printed, motorized blood smear device that prepares two smears simultaneously from a printed stage and a micro-motor drive with tunable linear velocity, controlled through a joystick-operated driver module. By spreading two slides in parallel with fully repeatable slide-to-slide motion, the device doubles throughput while eliminating operator-dependent motion artifacts, and can be fabricated on any benchtop 3D printer in under one day. To validate smear quality, we analyzed blood films from three donors together with a manual smear prepared by an expert from the blood of one of the same donors, giving a matched device-versus-manual pair. Automated Cellpose segmentation of 14,046 red blood cells across 12 bright-field fields showed that cell diameter was preserved and closely matched the expert smear (6.7-7.8 um across groups, within the 6.2-8.2 um human reference range; 6.7 vs 6.9 um in the matched pair), and that all films formed non-aggregated monolayers (Clark-Evans index of aggregation 1.04-1.24). Critically, red blood cells in the device films were markedly more circular than in the expert manual smear (mean eccentricity 0.405 vs 0.502; 0.443 vs 0.502 in the matched pair), with complete separation between the two methods at the level of whole fields of view. Because eccentricity reports smear-induced cell distortion, this indicates that a constant, mechanically controlled spreading velocity preserves red blood cell morphology better than skilled manual technique. BUsmear offers an accessible route to standardized smear geometry for quantitative analysis, including AI-based morphometry, and its low cost may be particularly advantageous in low-income countries with a high prevalence of malaria.

## Introduction

Blood smears remain central to hematological diagnosis, because many diseases manifest as characteristic changes in cell morphology. Red blood cells, the most abundant cells in blood, are anucleate biconcave discs of remarkably uniform size and shape, and this uniformity is what makes deviations from it diagnostically informative [1], [2]. Sickle cell disease [3], [4] produces sickled red blood cells; iron-deficiency anemia and thalassemia [5], [6] present with microcytosis and hypochromia; malaria [7]–[9] is diagnosed by directly visualizing intra-erythrocytic parasites; and various leukemias [10], [11] reveal blast cells and abnormal leukocyte distributions. Such morphology-based hallmarks can be recognized reliably only when smear quality is consistent, underscoring the need for standardized smear-preparation methods [12].

Conventional manual smear preparation remains highly operator-dependent [13], [14]: critical parameters such as spreading speed and initial droplet positioning must be adjusted intuitively by the user. The resulting films are frequently marred by surface contaminants and irregular distribution, yielding non-uniform layers that are too thick or too thin [15], [16]. Such inconsistencies and suboptimal sample placement directly degrade smear quality and, in turn, limit the reliability of downstream quantitative microscopy, machine-learning-based morphometry, and precise physical characterization of red blood cells. Therefore, there is a clear need for standardized mechanical intervention to ensure reproducible, high-quality cell monolayers.

Efforts to produce low-cost smear-preparation devices for non-expert users have accelerated in recent years, driven by advances in additive manufacturing such as 3D printing. Several groups have demonstrated automated and semi-manual smear devices built from 3D-printed components, showing that accessible fabrication can bring standardized smear preparation beyond specialized laboratories [17]– [20].

Building upon and extending these automated designs [17], [18], [20], we present automated blood smear device (BUs-mear), an original, low-cost, 3D-printed motorized smear platform. Unlike existing electronic systems that often require a continuous connection to a personal computer [17], [18], [20], our design adopts a fully standalone architecture built around an onboard Arduino Uno microcontroller. A key innovation is an analog joystick that gives the operator precise control over the initial position of the spreader slide relative to the blood droplet, allowing the pre-spread configuration to be fine-tuned without direct manual contact and readily accommodating different droplet volumes. The mechanical design is further optimized for high-throughput work by preparing two blood smears simultaneously under identical mechanical conditions: once the starting position is calibrated with the joystick, a constant smearing velocity is applied to both samples by the motorized control system. This dual-track approach increases efficiency while reducing manual variability, ensuring uniform monolayers suitable for high-fidelity biophysical profiling. To assess performance, we quantified the resulting films by automated cell segmentation and benchmarked their quality against expert-prepared and published automated smears.

## Materials and Methods

The machine consists of 3D-printed parts (Fig.1) and Commercial Off-The-Shelf (COTS) components. 3D-printed components form the main body of the machine while COTS components provide the mechanical motion and user-interface control. All the components used in the machine, are explained in the proceeding sections.

**Fig. 1.**
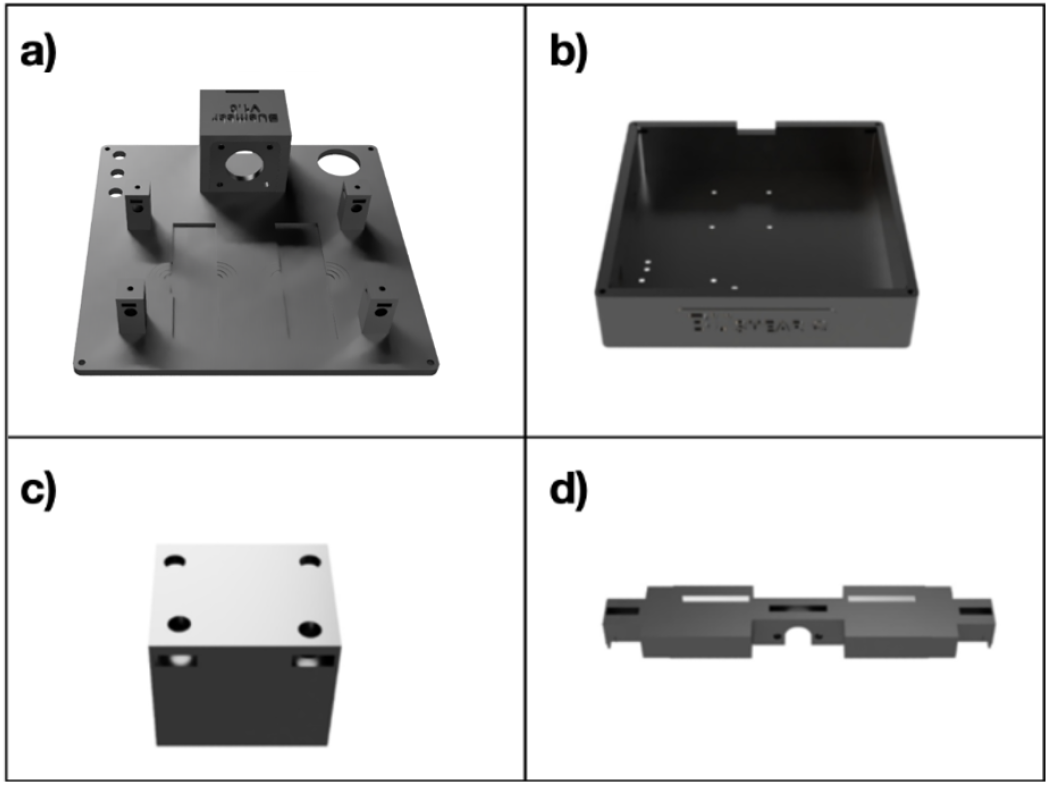
Four 3D-printed parts of the BUsmear device namely: a) the lid, b) the box, c) the joystick platform and d) the slider.

### 3D-printed Components

A Bambulab A1 printer with Acrylonitrile Butadiene Styrene (ABS) filament is used for printing the 3D-printed components of the device. The ABS filament is selected for its high tensile strength and durability. There are four 3D-printed parts which form the main body of the device given in Fig.1.

The components include the lid, which contains the housing for the stepper motor and, and the platform where smearing is performed. The box serves as a container for the Arduino Uno, stepper motor driver, and the joystick platform. The joystick platform is designed to elevate the joystick to the level of the lid. Lastly, the slider acts as a handle for the slider glass, moving linearly as the motor rotates to perform the smearing. Assembled device is demonstrated in Fig.2

**Fig. 2.**
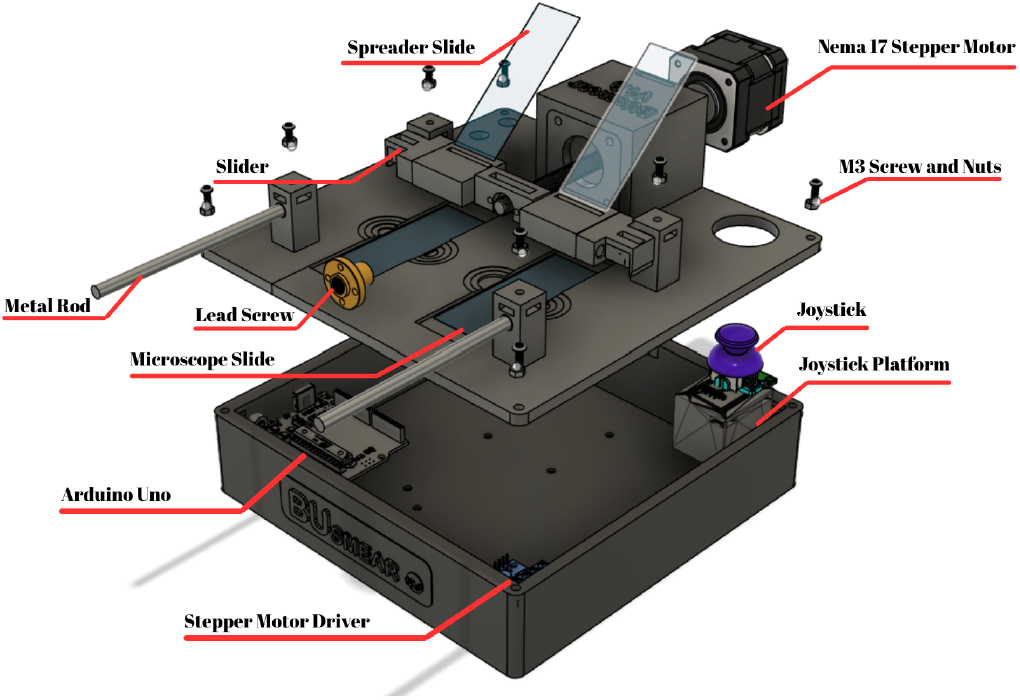
Labelled overview of the BUsmear device, showing the 3D-printed body and the electronic and mechanical components: the Arduino Uno and stepper-motor driver in the base; and the Nema 17 stepper motor, lead screw, metal-rod-guided slider, spreader slide, microscope slides, and joystick on the upper platform.

### COTS Components

COTS parts consists of electronic and non-electronic components. Electronic components create and control the motion while non-electronic components keep the device together, all of the COTS components are listed in Table I. The electronic components are connected as follows: The joystick and the motor driver is connected to the Arduino Uno, the motor driver is also connected to the motor so that it provides communication between the motor and the Arduino Uno. The motor driver and the Arduino Uno are connected to separate 7-12V power sources. The connection scheme is given in the Fig.3.

**TABLE 1.** List of components and their functions in the motion control system.

| Component | Function |
| --- | --- |
| Arduino Uno | Controls the electronic components |
| Nema 17 Stepper Motor | Provides rotational motion to drive the slider |
| Stepper Motor Driver | Enables communication between Arduino Uno and stepper motor (A4988) |
| Driver Expansion Board | Provides solder-free pin connections |
| Joystick | Allows user to control slider motion |
| Lead Screw | Converts rotational motion into linear motion |
| Rigid Coupler | Connects stepper motor to lead screw |
| Metal Rods | Provides sliding support for the slider |
| Screw and Nuts | M3 screws and nuts are used for assembling the components |

**Fig. 3.**
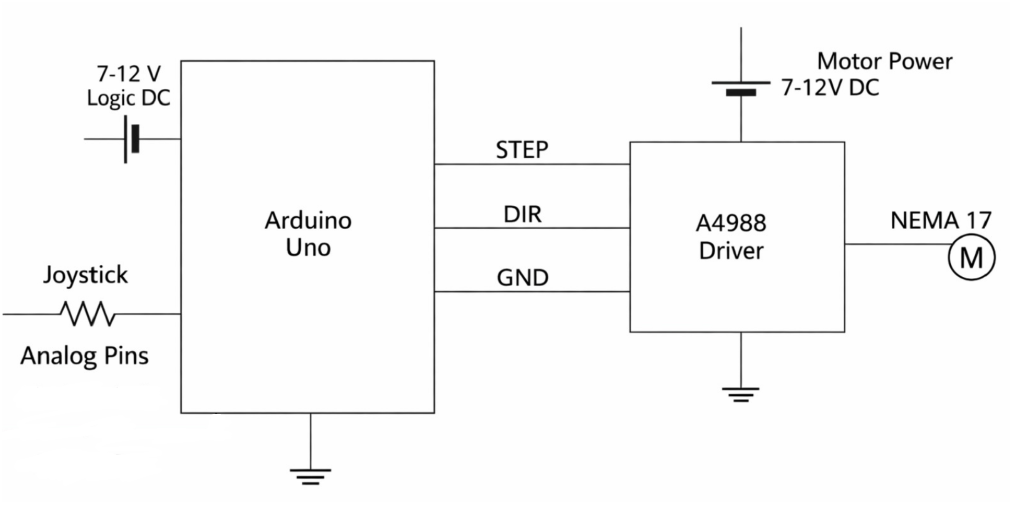
Schematic diagram of the BUsmear electronic control system, highlighting the independent power distribution and the interface between the Arduino Uno, the A4988 driver, and peripheral sensors.

### Electromechanical Implementation

User controls the motor which creates rotational motion, however linear motion is required to perform smear. The rotational motion of the motor is converted linear motion utilizing a lead screw shown in Fig.2. A lead screw nut is mounted on the lead screw and prevented from rotating by installing the slider on the lead screw nut. As the lead screw rotates, the lead screw nut moves on the lead screw, along with the slider. Spreader slides are placed in the slider and perform the blood smear as the slider moves. Real assembled BUsmear device is shown in Fig.4.

**Fig. 4.**
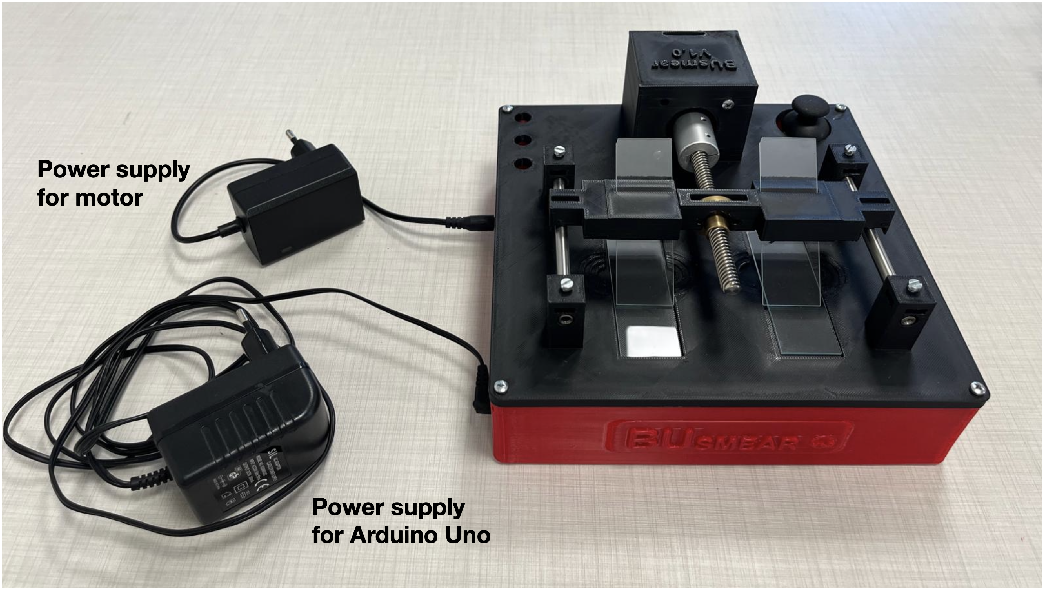
Overview of fully assembled BUsmear.

The code is uploaded to the Arduino Uno and orchestrates the operation of the electronic components in harmony. The user can adjust the motor speed by modifying the parameter period in the code, which defines the period of the motor steps. The motor’s motion is governed by the joystick input. Rotation begins when the joystick is fully pushed or pulled, and continues until the joystick is released beyond the halfway position. The electronic control system is housed within the 3D-printed base of the device, as shown in the open-view assembly in Fig. 5.

**Fig. 5.**
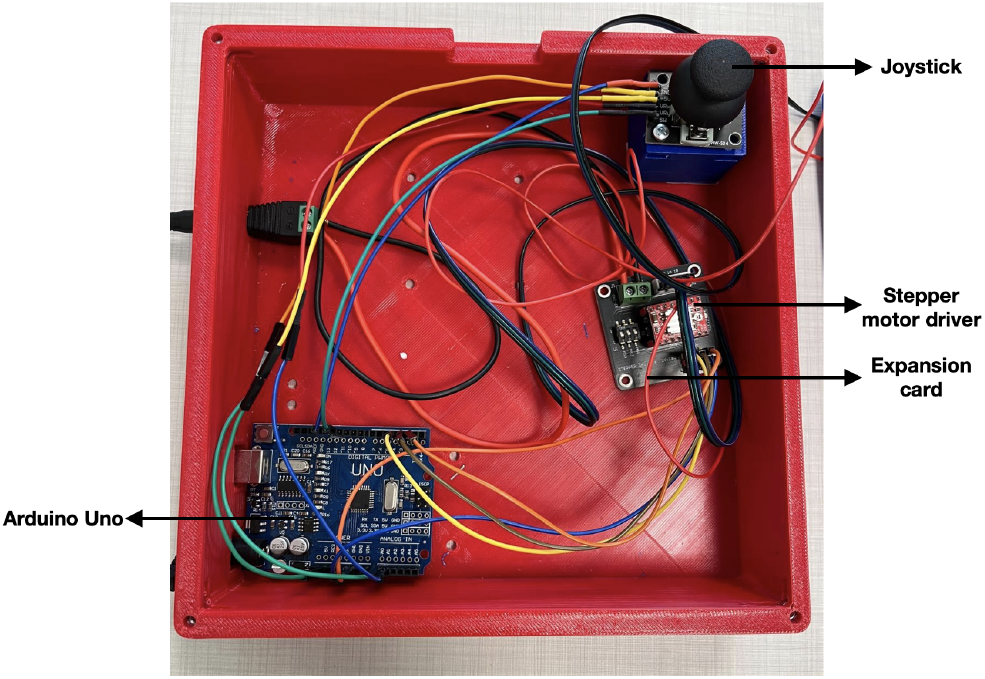
Open-view of the controlbox of BUsmear.

An Arduino Uno microcontroller serves as the central processing unit, executing the motion control algorithms. The user interface is provided through an integrated analog joystick, enabling manual positioning and calibration of the spreader mechanism. To ensure precise and repeatable linear motion, a high-resolution stepper motor driver is employed, mounted on a dedicated expansion board for simplified wiring and power distribution. This compact integration of electronics within the structural frame enhances both portability and stability of the system during laboratory use.

## Results

### Automated image analysis of BUsmear films

Blood smears prepared with BUsmear were imaged unstained in bright field (Zeiss PALM, LD Plan-Neofluar 20 *×* /0.4; field of view 674.8 *×* 510.9 *µ*m, 0.349 *µ*m px^*™*1^).Three smears were prepared from 3 donors using BUsmear device. As a control, an expert prepared a manual smear from one of the donor. Twelve fields (three per group) were segmented with Cellpose [23], [24], yielding 14 046 red blood cells from which cell count, surface density, diameter, eccentricity and two spatial-distribution metrics were computed (Fig. 6, Table II). red blood cells were detected reliably across the full density range spanned by the films, from 1.6 to 7.4 *×* 10^3^ cells mm^*™*2^ (Fig. 6a–d), with the automated masks closely following individual cell outlines (Fig. 6e).

**TABLE 2.**
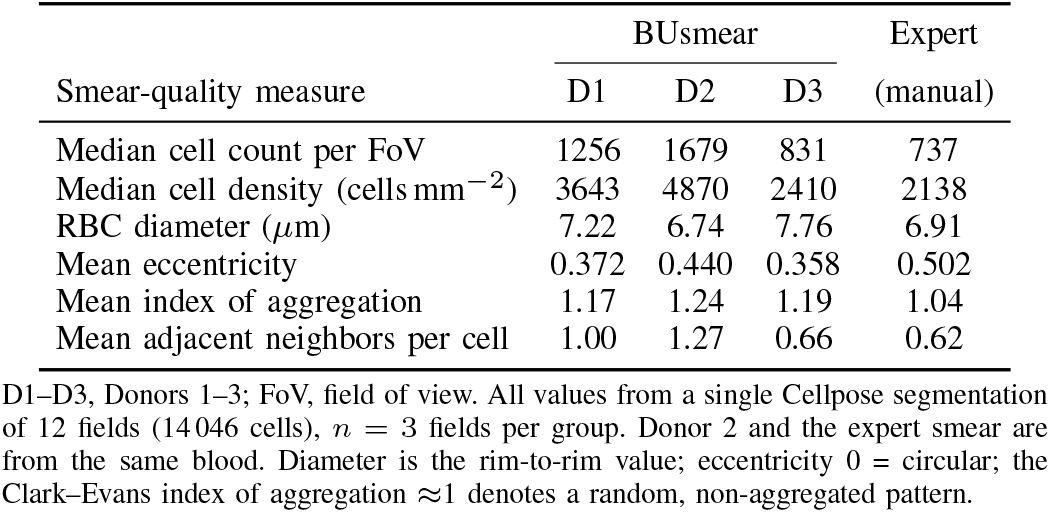
Smear-Quality Metrics for Three BUsmear Films and an Expert-Prepared Manual Smear.

**Fig. 6.**
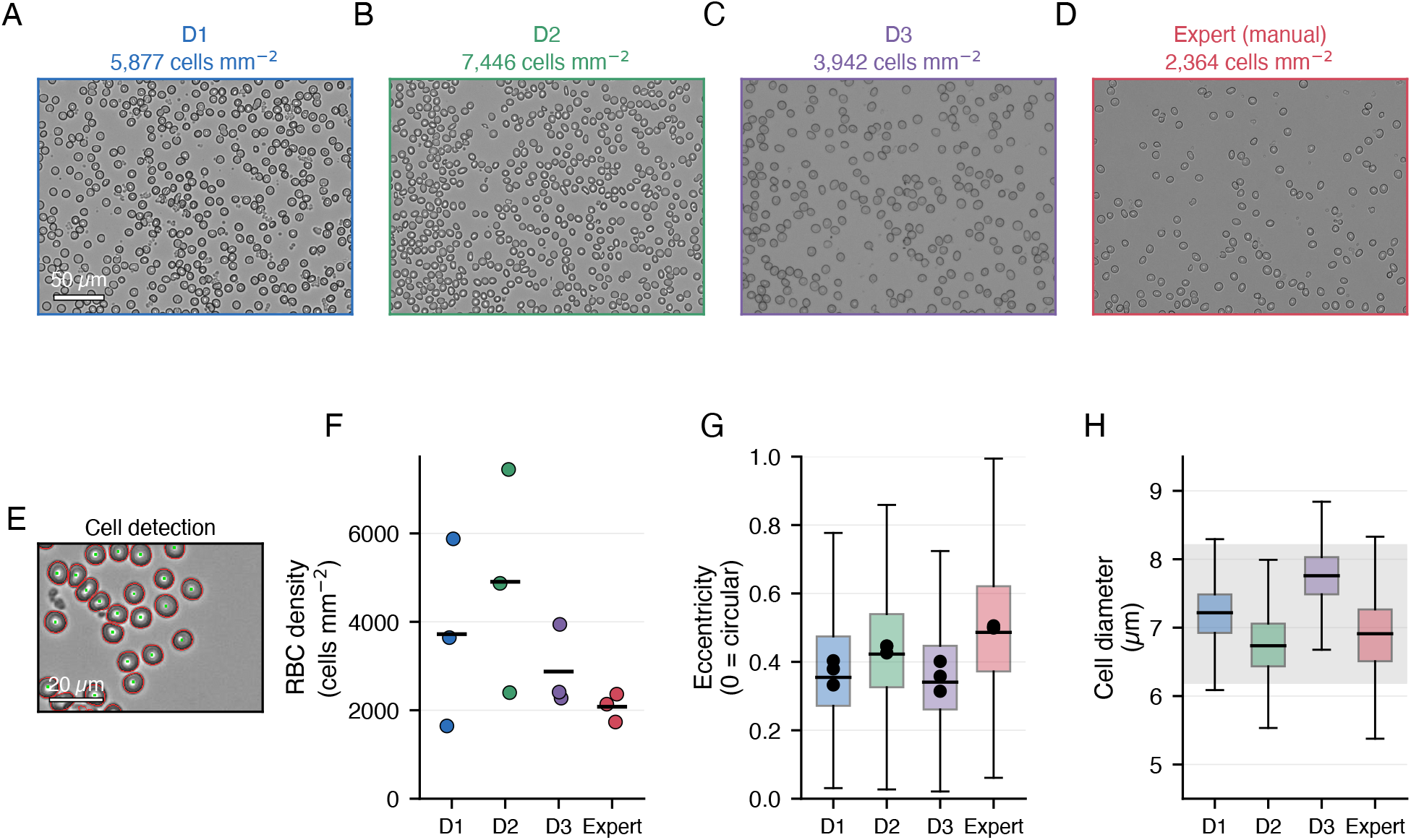
BUsmear films from three donors compared with an expert-prepared manual smear. (**a–d**) Representative bright-field fields for Donors 1–3 (D1–D3, BUsmear) and the expert manual smear; for each group the sharpest of its three fields is shown, and the quoted density is that of the displayed field. Scale bar (a), 50 *µ*m.(**e**) Automated cell detection: Cellpose outlines (red) and centroids (green) of the red blood cells entering the analysis. Scale bar, 20 *µ*m. (**f**) RBC surface density; points, individual fields (*n* = 3 per group); bars, group mean. (**g**) Cell eccentricity (0 = circular). Boxes, per-cell distribution (median, IQR, 1.5 *×* IQR whiskers); black points, per-field means. red blood cells in the device films are more circular than in the expert manual smear (pooled 0.405 vs 0.502; Mann–Whitney *p* < 0.001, Cohen’s *d* = 0.57). (**h**) Rim-to-rim cell diameter; grey band, reference human-red blood cell range (6.2–8.2 *µ*m) [21], [22]. Donor 2 and the expert smear were prepared from the same blood, giving a matched device-versus-manual pair. All metrics derive from a single Cellpose segmentation of 12 fields (14 046 cells); diameters from an independent rim-to-rim measurement (*n* = 6127 cells).

### Cell size is preserved and equivalent to the manual smear

The diameter of detected red blood cells, measured rim-to-rim independently of the Cellpose masks (*n* = 6127 cells), fell within the reference range for human red blood cells (6.2–8.2 *µ*m) for every group: 7.22, 6.74 and 7.76 *µ*m for Donors 1–3 and 6.91 *µ*m for the expert smear (Fig. 6h, Table II). Between-donor differences (6.7–7.8 *µ*m) are consistent with normal inter-individual variation in red blood cell size. Crucially, in the matched-blood pair the device and the expert gave essentially the same diameter (Donor 2, 6.74 *µ*m vs expert, 6.91 *µ*m), showing that mechanised spreading does not alter cell size relative to expert manual technique.

### sBUsmear preserves cell shape better than the manual smear

Cell shape, by contrast, differed clearly between the two preparation methods. red blood cells in BUsmear films were more circular than in the expert’s manual smear: mean eccentricity 0.372, 0.440 and 0.358 for Donors 1–3 versus 0.502 for the expert (0 = circular; Fig. 6g). Pooling the device films (*n* = 11 896 cells) gives 0.405 versus 0.502 for the expert (*n* = 2150 cells; Mann–Whitney *p* < 0.001, Cohen’s *d* = 0.57). The effect persists in the matched-blood comparison, where donor identity cannot contribute: Donor 2 versus the expert smear from the same blood gave 0.443 versus 0.502 (*p* < 0.001, *d* = 0.35).

Because the relevant unit of comparison is the smear rather than the individual cell, we also assessed the effect at the level of whole fields of view. Here the separation is complete: the nine device fields spanned mean eccentricities of 0.314–0.446, whereas the three expert fields fell at 0.499, 0.501 and 0.505, so that the most elongated device field was still less eccentric than the least eccentric expert field (Welch’s *t*-test, *p* < 0.001; Fig. 6g). Averaging the *∼*1000 cells in each field cancels cell-to-cell variability and isolates the smear-level signal, which is why the per-cell distributions overlap appreciably (a randomly chosen expert cell is more elongated than a randomly chosen device cell in 66% of comparisons) while the field means do not overlap at all. Single cells therefore cannot be assigned to a preparation method, but smears can.

Importantly, this difference is not a density artefact. Within the device films eccentricity increased with density (Donor 3, 2410 cells mm^*™*2^, 0.358; Donor 1, 3643, 0.372; Donor 2, 4870, 0.440), yet the expert smear had both the *lowest* density (2138 cells mm^*™*2^) and the *highest* eccentricity (0.502). Density therefore acts against the observed difference, implying that the true shape advantage of the device is at least as large as measured.

### Spatial distribution

All films were well dispersed rather than clumped, with Clark–Evans indices of aggregation between 1.04 and 1.24 (a value of 1 denotes a random pattern; Table II). The expert smear was closest to random (1.04) and the device films slightly more dispersed (1.17–1.24). Mean touching neighbours per cell was higher in the device films (0.66–1.27) than in the expert smear (0.62), consistent with their higher cell density; the density-independent index of aggregation shows that none of the films were aggregated.

### Benchmarking against published values

To place these measurements in context, we compared them with the values reported for Autohaem [17] (Table III). The eccentricity we measured for our own expert (0.502) is almost identical to the mean reported for expert operators in that study (0.498), despite different instruments, laboratories and blood samples—an external check that the segmentation and shape measurement used here are comparable to the published pipeline [17]. Against this reference, BUsmear films remain the more circular (0.405), and their index of aggregation (1.20) is close to that of expert operators (1.11) and of the optimised Autohaem devices (1.09–1.17). Median density of the device films (3.6 *×* 10^3^ cells mm^*™*2^) exceeds that of expert operators and of Autohaem smear+ (2.5–3.0 *×* 10^3^ cells mm^*™*2^).

**TABLE 3.** BUsmear and Expert Control Benchmarked Against the Conditions Reported for Autohaem [17].

| Smear-quality measure | This work |  | Autohaem |  |  |
| --- | --- | --- | --- | --- | --- |
|  | BUSmear<br>45° | expert<br>manual | experts<br>~45° | smear<br>30° | smear+<br>50° |
| Median density (cells mm<sup>-2</sup>) | 3643 | 2138 | 2528 | 2455 | 3023 |
| Index of aggregation | 1.20 | 1.04 | 1.11 | 1.09 | 1.17 |
| Mean eccentricity | 0.405 | 0.502 | 0.498 | 0.535 | 0.469 |
| Adjacent neighbors* | 0.97 | 0.62 | 0.17 | 0.33 | 0.20 |

Eccentricity 0 = circular; index of aggregation $\approx 1$ denotes a random, non-aggregated pattern. The Autohaem smear+ condition is its optimum, 50° at 7 cm s<sup>-1</sup>. Autohaem counts were converted to density using their 188 × 117 μm field of view. BUSmear values summarize the three donor films (nine fields): the density is the median across fields, the index of aggregation and adjacent-neighbor counts are means across fields, and the eccentricity is the cell-weighted mean over all 11 896 segmented cells, as in the Results. The unweighted field-level mean eccentricity is 0.390, so the value quoted here is the more conservative of the two.
\* Adjacent-neighbor counts are not comparable across pipelines: our expert smear (0.62) and the Autohaem experts (0.17) differ by more than threefold although both are expert-prepared manual smears, reflecting different touching-tolerance definitions. The measure also scales with cell density. The density-independent index of aggregation ( $\approx 1$ ) shows that none of the films were aggregated.

### Blood sampling and smear preparation

Capillary blood was obtained from the fingertip of each participant with a sterile lancet. Smears were prepared from three donors using BUsmear, with the spreader slide held at a fixed 45^*?*^ angle, the clinically recommended manual spreading angle [25], and driven at a constant linear velocity of 2.5 cm s^*™*1^. A 2 *µ*L droplet was deposited on the slide and the spreader was brought to its starting position with the joystick before each run. As a control, an expert operator prepared a manual smear from his own blood using the conventional wedge technique; because that donor’s blood was also processed with BUsmear (Donor 2), the data set contains a matched device-versus-manual pair in which only the preparation method differs. Two smears were prepared per donor. All films were air-dried for 2 min and imaged unstained, without fixation.

### Microscopy

Films were imaged in transmitted bright field on a Zeiss PALM system with an LD Plan-Neofluar 20 *×* /0.4 Korr M27 objective. Three fields were acquired per group (twelve in total), selected by the operator within the monolayer region of each film.

### Image segmentation and cell metrics

Images were converted to 8-bit greyscale and segmented with Cellpose 3.1.1 [23] using the generalist `cyto3` model with a cell-diameter prior of 22 px (*≈*7.7 *µ*m), single-channel input and `flow_threshold` = 0. Masks with an area outside 90–1500 px were discarded as debris or merged objects; all reported quantities derive from the retained masks of this single segmentation.

For every retained cell we computed the area, the centroid and the second central moments of the mask. *Cell count* is the number of retained masks and *surface density* the count divided by the field area. *Eccentricity* was obtained from the second moments as 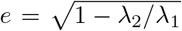, where *λ*_1_ and *λ*_2_ are the major and minor eigenvalues of the moment matrix, so that *e* = 0 describes a circle and *e* = 1 a line; this is the definition used by CellProfiler and by the reference study [17].

Spatial organisation was quantified by two measures. The *Clark–Evans index of aggregation* [26] is *R* = *r*_obs_*/r*_exp_, where *r*_obs_ is the mean nearest-neighbour centroid distance over all cells not touching the image border and 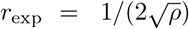 is the value expected for a Poisson pattern of the same intensity *ρ* = *N/A*; *R ≈* 1 indicates a random pattern, *R* < 1 clustering and *R >* 1 dispersion. *Adjacent neighbours* counts touching cells: labels were expanded by up to 2 px into the background to bridge the thin gaps left by segmentation, and two cells were scored as adjacent where their expanded regions became 4-connected; the reported value is the mean number of distinct neighbours per cell.

Cell diameter was measured independently of the Cellpose masks, because a mask follows the outer edge of the bright-field refraction ring and therefore overestimates the cell by *≈*0.8 *µ*m. Candidate cells were detected by local-background subtraction (61-px uniform filter, Gaussian smoothing *σ* = 1 px) followed by thresholding and morphological cleaning, and only round, solid, isolated objects were retained (area 120–900 px, bounding-box aspect ratio 0.75–1.33, fill fraction *>*0.62). For each retained cell the mean radial intensity was sampled along 72 directions at radii of 2–18 px in 0.5-px steps; the rim-to-rim diameter was taken as twice the radius of the intensity minimum, the darkest point of the refraction ring, which marks the cell edge, refined to sub-pixel accuracy by parabolic interpolation. Values outside 4–13 *µ*m were rejected.

### Statistical analysis

Eccentricity distributions were compared at two levels. At the level of individual cells, groups were compared with two-sided Mann–Whitney *U* tests and the effect size reported as Cohen’s *d* using the pooled standard deviation; the probability that a randomly chosen cell from one group exceeds a randomly chosen cell from the other was obtained as Φ(*d/* 2).

Because the unit of comparison relevant to smear quality is the film rather than the cell, group means were also compared at the level of whole fields of view using Welch’s unequal-variance *t*-test on the per-field mean eccentricity. All analyses were performed in Python 3 with NumPy, SciPy and pandas; segmentation used Cellpose 3.1.1 on PyTorch 2.2.

## Discussion

We evaluated the blood smears produced by BUsmear, a dual-smear device with joystick control that builds on the autohaem smear+ platform [17], by automated image analysis of films from three donors, benchmarked against a manual smear prepared by an expert. Because the expert smeared his own blood, which was also processed with the device, the dataset contains a matched pair in which donor identity is held constant and only the preparation method differs. Across 14 046 red blood cells segmented from twelve bright-field fields with Cellpose [23], the device produced films that were correctly sized, non-aggregated and, most notably, better preserved in shape than the expert’s manual smear.

The central finding is that red blood cells in BUsmear films are more circular than those in the manual smear (mean eccentricity 0.405 vs 0.502 pooled, and 0.443 vs 0.502 in the matched-blood pair). Eccentricity is the established readout of smear-induced cell distortion: poorly executed spreading, for example pushing rather than pulling the blood, elongates red blood cells [17]. That a mechanised device outperforms a trained operator on this measure is consistent with its defining property, a constant, reproducible spreading velocity, removing the acceleration and hesitation inherent to hand-drawn smears. The difference is unlikely to be an artefact of cell density: within the device films eccentricity rose with density, yet the expert smear combined the lowest density with the highest eccentricity, so the density gradient works against the observed effect rather than producing it. It is also robust to the level of analysis. Although the per-cell distributions overlap appreciably (Cohen’s *d* = 0.57), averaging the *∼*1000 cells in each field removes cell-to-cell variability and separates the two methods completely, with every device field less eccentric than every expert field. As an external check on the measurement itself, the expert value we obtained (0.502) is almost identical to the mean eccentricity reported for expert operators in the autohaem study (0.498) [17], despite different instruments and laboratories, supporting the comparability of our pipeline.

Cell size, by contrast, was unaffected by the preparation method: in the matched-blood pair the device and the expert gave 6.7 and 6.9 *µ*m respectively, and all four groups fell within 6.7–7.8 *µ*m, inside the human-red blood cell reference range. This is the expected result, spreading should not change how large a cell is, and it serves as a positive control confirming that the device introduces no size bias. All films were likewise well dispersed (Clark–Evans index of aggregation 1.04–1.24), so the shape advantage is not accompanied by any penalty in cell distribution. Together, the size, shape and spatial results indicate that BUsmear yields monolayers at least as faithful as expert-prepared films while removing the operator dependence that motivated the device; combined with simultaneous dual-smear preparation this doubles throughput without a quality penalty, and the fully 3D-printed, standalone construction keeps the platform inexpensive and independent of a host computer.

Mean touching-neighbour counts were higher in the device films (0.97) than in the expert smear (0.62), but this measure scales with cell density, which was higher in the device films, and is not comparable across analysis pipelines: our expert smear and the Autohaem experts differ more than threefold (0.62 vs 0.17) although both are expert-prepared manual smears. The density-independent index of aggregation confirms that none of the films were clumped. Finally, all films were imaged on one microscope with a single segmentation pipeline; this makes the internal comparison tightly controlled, but absolute values may shift with other imaging systems. As the Autohaem study reports no exact 45^*?*^ condition, our benchmarking against it brackets BUsmear’s fixed 45^*?*^ angle, the clinically recommended manual spreading angle, between the nearest published settings (expert operators at *∼*45^*?*^ and the smear+ optimum at 50^*?*^). Future work will broaden the donor and operator panels and integrate imaging and counting directly into the device.

## Conclusion

We have presented BUsmear, a low-cost, fully 3D-printed blood-smear device that prepares two smears simultaneously under joystick control from a standalone Arduino Uno-based drive. Smear quality was assessed by automated Cellpose-based analysis of 14 046 red blood cells from three device-prepared smears, benchmarked against a manual smear prepared by an expert from the blood of one of the same donors. Cells were correctly sized, with diameters of 6.7–7.8 *µ*m across all groups, inside the 6.2–8.2 *µ*m human-red blood cell reference range [21] and closely matching the expert smear prepared from the same blood (6.74 vs 6.91 *µ*m), confirming that mechanised spreading introduces no size bias. All films were well dispersed and non-aggregated (Clark–Evans index of aggregation 1.04–1.24). Critically, the red blood cells were more circular than in the manual smear (mean eccentricity 0.405 vs 0.502, and 0.443 vs 0.502 in the matched-blood pair), with the two methods separating completely at the level of whole fields of view.

Because eccentricity reports smear-induced cell distortion, this result indicates that a constant, mechanically controlled spreading velocity preserves red blood cell morphology better than skilled manual technique, while simultaneous dual-smear preparation doubles throughput at no cost to quality. BUsmear therefore offers a practical and inexpensive route to standardised blood films, particularly where trained operators or commercial smearing instruments are scarce. In future work, we aim to integrate an onboard microscope and automated cell-counting software into the platform, yielding a single, self-contained device that prepares, images and quantifies blood smears without external instrumentation, and to extend the validation to larger donor and operator panels.

## Ethical Statement

The experimental protocols, including blood sample, were approved by Boğaziçi University Science and Engineering Fields Human Research Ethics Committee (FMINAREK) with the ethical permission number 2025-15. Blood samples were taken using a lancet needle from the participants’ fingertips. Informed consent was obtained from all participants. All methods were performed in accordance with Declaration of Helsinki.

## Supplementary Materials

The design and video of the BUsmear can be found in Gitlab.

## Author Contributions

T.P. and B.A. initiated the study. T.P., M.T.K., and B.C.E. designed the device. M.T.K. assembled the electronic components. T.P. and M.T.K. performed the analysis. T.P., M.T.K. and B.C.E. wrote the manuscript. All authors read and approved the final manuscript.

